# Automatization and validation of the hippocampal-to-ventricle ratio in a clinical sample

**DOI:** 10.1101/2024.04.12.588928

**Authors:** Sofia Fernandez-Lozano, Vladimir Fonov, Dorothee Schoemaker, Jens Pruessner, Olivier Potvin, Simon Duchesne, D. Louis Collins, Alzheimer’s Disease Neuroimaging Initiative

**Affiliations:** McConnell Brain Imaging Centre, Montreal Neurological Institute, McGill University, Montreal, Quebec, Canada; Department of Neurology and Neurosurgery, McGill University, Montreal, Quebec, Canada; Department of Psychology, McGill University, Montreal, Quebec, Canada.; Department of Psychology, University of Konstanz, Konstanz, Germany.; Quebec Heart and Lung Research Institute, Quebec City, Quebec, Canada; Radiology and Nuclear medicine Department, Université Laval, Quebec City, Quebec, Canada; Department of Biomedical Engineering, McGill University, Montreal, Quebec, Canada

**Keywords:** MRI, Hippocampus, Alzheimer’s Disease, Automatic segmentation, Memory, Cognitive decline

## Abstract

**Background:** The hippocampal-to-ventricle ratio (HVR) is a biomarker of medial temporal atrophy, particularly useful in the assessment of neurodegeneration in diseases such as Alzheimer’s disease (AD). To minimize subjectivity and inter-rater variability, an automated, accurate, precise, and reliable segmentation technique for the hippocampus (HC) and surrounding cerebro-spinal fluid (CSF) filled spaces — such as the temporal horns of the lateral ventricles — is essential.

**Methods:** We trained and evaluated three automated methods for the segmentation of both HC and CSF (Multi-Atlas Label Fusion (MALF), Nonlinear Patch-Based Segmentation (NLPB), and a Convolutional Neural Network (CNN)). We then evaluated these methods, including the widely used *FreeSurfer* technique, using baseline T1w MRIs of 1,641 participants from the AD Neuroimaging Initiative study with various degree of atrophy associated with their cognitive status on the spectrum from cognitively healthy to clinically probable AD. Our gold standard consisted in manual segmentation of HC and CSF from 80 cognitively healthy individuals. We calculated HC volumes and HVR and compared all methods in terms of segmentation reliability, similarity across methods, sensitivity in detecting between-group differences and associations with age, scores of the learning subtest of the Rey Auditory Verbal Learning Test (RAVLT) and the Alzheimer’s Disease Assessment Scale 13 (ADAS13) scores.

**Results:** Cross validation demonstrated that the CNN method yielded more accurate HC and CSF segmentations when compared to MALF and NLPB, demonstrating higher volumetric overlap (Dice Kappa = 0.94) and correlation (rho = 0.99) with the manual labels. It was also the most reliable method in clinical data application, showing minimal failures. Our comparisons yielded high correlations between FreeSurfer, CNN and NLPB volumetric values. HVR yielded higher control:AD effect sizes than HC volumes among all segmentation methods, reinforcing the significance of HVR in clinical distinction.

**Associations:** The positive association with age was significantly stronger for HVR compared to HC volumes on all methods except FreeSurfer. Memory associations with HC volumes or HVR were only significant for individuals with mild cognitive impairment. Finally, the HC volumes and HVR showed comparable negative associations with ADAS13, particularly in the mild cognitive impairment cohort.

**Conclusion:** This study provides an evaluation of automated segmentation methods centered to estimate HVR, emphasizing the superior performance of a CNN-based algorithm. The findings underscore the pivotal role of accurate segmentation in HVR calculations for precise clinical applications, contributing valuable insights into medial temporal lobe atrophy in neurodegenerative disorders, especially AD.

**Authorship:**

- Sofia Fernandez-Lozano: Conceptualization, Methodology, Software, Investigation, Writing – Original Draft, Visualization.
- Vladimir Fonov: Software, Data Curation.
- Dorothee Schoemaker: Resources, Writing – Review & Editing.
- Jens Pruessner: Resources, Writing – Review & Editing.
- Olivier Potvin: Resources, Writing – Review & Editing.
- Simon Duchesne: Resources, Writing – Review & Editing.
- D. Louis Collins: Conceptualization, Writing – Review & Editing, Supervision.

## 1. Introduction

The hippocampus (HC) is a medial temporal lobe structure located medially to the lateral horn of the lateral ventricles, posterior to the amygdala, and superior to the parahippocampal gyrus. Integral to the limbic system, the HC plays an important role in cognition, more specifically for declarative and semantic memory (Eichenbaum 2004). It is further implicated in a number of psychiatric disorders such as depression (Campbell et al. 2004; Videbech and Ravnkilde 2004), schizophrenia (Adriano, Caltagirone, and Spalletta 2012), and addiction (Morimoto et al. 2018), as well as neurological diseases such as Alzheimer’s disease (AD) (Jack et al. 2018), Parkinson’s disease (Kandiah et al. 2014), multiple sclerosis (Koenig et al. 2014), or temporal lobe epilepsy (Reyes et al. 2018).

The HC is affected by normal and pathological effects related to aging. Its atrophy, measured by structural MRI, is one of the best documented pathological features in AD (Jack et al. 2018). The yearly rate of hippocampal volume reduction is reported as 1.4% in healthy populations and up to 4.7% in dementia due to AD (Barnes et al. 2009). This disparity in the rate of atrophy between cognitively healthy controls, individuals with mild cognitive impairment (MCI), a prodromal stage of AD, or patients with AD has been widely reported in other cross-sectional and longitudinal studies (Frankó, Joly, and Alzheimer’s Disease Neuroimaging Initiative 2013; Henneman et al. 2009; Shi et al. 2009). The rate of HC atrophy may even help to distinguish patients with cognitive impairment who progress to dementia from those who do not (Coupé et al. 2011; Zandifar et al. 2017).

Although manual segmentation is considered the gold-standard in the volumetric assessment of the HC, it suffers from several drawbacks that impede its practical application in large datasets. Despite the availability of software tools with semi-automated guidance and/or 3D visualization, manual segmentation of the HC is a laborious and time-consuming task that can take up to two hours per subject (Schoemaker et al. 2019). It is affected by high intra- and inter-rater variability, even in experts (Dill, Franco, and Pinho 2015). As a response to these limitations, several automated segmentation methods have been proposed; for reviews see (Dill, Franco, and Pinho 2015; Yi et al. 2021).

The most common methods are based on label propagation, or segmentation using one or multiple templates, given their efficacy and accuracy. In a performance comparison done by Zandifar et al (2017) on four different methods (ANIMAL with template library and label fusion (Collins and Pruessner 2010), nonlocal patch-based segmentation with expert priors (Coupé et al. 2011; Fonov et al. 2012), and FreeSurfer v5.3 (Fischl et al. 2002), the nonlocal patch-based method was shown as being the most accurate, based on overlap statistics with manually labeled ground truth data.

The explosive development of deep learning has led to new methods and strategies for segmentation. Amongst the most successful, the U-net (Ronneberger, Fischer, and Brox 2015) is a Convolutional Neural Network (CNN) with a U-shape network structure as its name indicates. It is characterized by a contraction path that captures context information while down-sampling the image and an expansion path that allows precise positioning on which image up-sampling and feature map fusion are repeated. It has shown to segment HC with Dice scores ranging between 0.89 and 0.92 (Yi et al. 2021).

Even if one had access to the most accurate segmentation method, volumetric studies of the HC show limitations due to the structure’s high inter-subject variability in shape and volume, even in cognitively healthy populations (Lupien et al. 2007; Nobis et al. 2019). For this reason, researchers have explored different strategies to improve the reliability and robustness of the diagnostic or prognostic impact of HC volumetry. One such strategy is to normalize HC volume with the volume of the cerebrum. Using the HC/cerebrum ratio, investigators were able to successfully identify different AD subtypes based on the spatial distribution of tau pathology, as well as predict faster cognitive decline (Risacher et al. 2017; Whitwell et al. 2012).

In addition to cortical atrophy, ventricular expansion is commonly present in neurodegenerative disorders, in particular in MCI and AD (Apostolova et al. 2006; 2013; Dalaker et al. 2011; Mak et al. 2017; Seif, Ziegler, and Freund 2018). Leveraging the idea of ex-vacuo dilation, the HC occupancy is a single biomarker of medial temporal atrophy consisting of the ratio of the volume of the HC to the sum of the volumes of the HC and inferior lateral ventricle (Heister et al. 2011). The median survival times in relation to conversion from MCI to AD were found to be significantly shorter for those subjects at risk due to atrophy measured by low HC occupancy than those at risk from any other measure or combination of measures from learning performance, CSF tau, amyloid and tau/amyloid ratio.

The HC-to-ventricle ratio (HVR) as an extension of the original idea of a ratio combining in a single metric the estimation of HC volume with the surrounding ventricular enlargement in a similar manner to HC occupancy, but instead using the complete lateral horn of the ventricle (Schoemaker et al. 2019). When calculated in two cohorts of cognitively normal aging adults, the HVR showed stronger relationships to age and delayed memory than the raw hippocampal volumetric measure.

The goal of this paper was to apply and compare available automatic segmentation techniques to automate HVR estimation (Pruessner et al. 2000; Schoemaker et al. 2019), explore potential variations introduced by the different segmentation methods, and then to use this automatic method to replicate and extend the earlier findings in a larger sample of AD, MCI and cognitively healthy individuals.

## 2. Materials and methods

### 2.1. Data

The data used in the preparation of this article was obtained from the Alzheimer’s Disease Neuroimaging Initiative (ADNI) database (https://adni.loni.usc.edu). ADNI was launched in 2003 as a public-private partnership, led by Principal Investigator Michael W. Weiner, MD. The primary goal of ADNI has been to test whether serial magnetic resonance imaging (MRI), positron emission tomography, other biological markers, and clinical and neuropsychological assessment can be combined to measure the progression of MCI and early AD. Specifically for this study, we used baseline 3D T1w MRIs from the ADNI-1, ADNI-2, and ADNI-GO cohorts.

Training of the segmentation algorithms was done using manual HC labels and the surrounding temporal horns of the lateral ventricles (N = 80). This data and the manual segmentation protocol are identical to those described in (Schoemaker et al. 2019). The labels were segmented by trained neuroanatomical experts and are considered as the ground-truth for HC volumes (HCvol) and perihippocampal ventricular space (CSFvol).

Demographic, memory, and cognitive data for the included subjects were downloaded from the ADNI website. To assess memory, we used the scores from the Learning subtest of the Rey Auditory Verbal Learning Test (RAVLT). As a measure of global cognitive function, we used scores from the AD Assessment Scale-13 (ADAS-13).

### 2.2. Image preprocessing

Preprocessing consisted of denoising (Coupe et al. 2008), N3 image intensity inhomogeneity correction (Sled, Zijdenbos, and Evans 1998), linear intensity normalization based on histogram matching between the image and the average template, and affine registration to a stereotaxic space defined as a population-specific template created from the ADNI-1 cohort (Fonov et al. 2011).

### 2.3. Automated segmentation methods

We trained three automatic segmentations methods described below using the HC and CSF manual labels. Additionally, we compared our trained methods with the output given by different versions of FreeSurfer with its preprocessing and processing pipelines (Fischl et al. 2002).

#### 2.3.1. Multi-Atlas Label Fusion

The first segmentation method consisted of a multi-atlas label fusion (MALF) approach. Using an augmented ANIMAL segmentation method (Collins et al. 1995; Collins and Evans 1997), this algorithm is enhanced with the use of a template library and label fusion (Collins and Pruessner 2010). Training consisted in a Monte Carlo Cross-Validation strategy (Shao 1993), that is, for every potential segmentation a template library was built from a random subset of the manual labels (n=64).

#### 2.3.2. Nonlinear Patch-Based Segmentation

The second segmentation method was the Nonlinear Patch-Based Segmentation (NLPB) with a population-specific template (Coupé et al. 2011; Fonov et al. 2012; H. Wang et al. 2013). Similar to MALF, this algorithm was trained using Monte Carlo Cross-Validation where a template library was created from a random subset of the manual labels (n=64). Additionally, manual labels were non-linearly aligned to a template space constructed from the ADNI-1 database (Fonov et al. 2011).

#### 2.3.3. Convolutional Neural Network

Lastly, we trained a deep learning CNN based on a 3D version of the U-Net architecture (Ronneberger, Fischer, and Brox 2015). It uses a combination of the Dice kappa overlap and a multi-label Hausdorff-like distance approximation as loss function. Further details were reported previously (Fonov, Rosa-Neto, and Collins 2022). To train the network, we used a 5-Fold cross validation strategy (i.e., split the data into a training set of 64 images with their manual labels and a validation set consisting of the remaining unlabeled 16 images). The training data was augmented with geometric transformations, random signal shift, amplification, and addition of voxel-level additive noise by a factor of 16 before training the network for 200 epochs for each cross-validation split.

#### 2.3.4. FreeSurfer

We compared the output of our trained segmentations with those obtained from FreeSurfer (https://surfer.nmr.mgh.harvard.edu). First, we used the values of HCvol from UCSF from either version 4.3 (N=545) or version 5.1 (N=701) of the cross-sectional FreeSurfer pipeline reported by ADNI on the files UCSFFSX_11_02_15.csv and UCSFFSX41_11_08_19.csv respectively, and our own results using cross-sectional FreeSurfer pipeline version 6.0 with default parameters. Since ADNI does not report inferior lateral ventricle volumes for the earlier FreeSurfer results, we calculated HVR using only FreeSurfer version 6.0.

### 2.4. Hippocampal measures

Using the output from the automatic segmentations we obtained HCvol and calculated HVR for each of the different methods. HVR was calculated using HCvol and CSFvol:

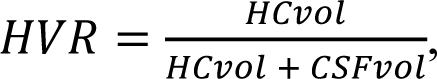

When calculating HVR using the FreeSurfer volumes, CSFvol was defined as the volume of the inferior lateral ventricle. For both HCvol and HVR, we use the mean of both sides in all our analyses.

We also normalized HCvol by head size for each technique, using the same volumetric scale factor obtained from the affine registration to stereotaxic space.

### 2.5. Segmentation performance assessment

#### 2.5.1. Cross-validation

After training the automatic segmentation methods using manual labels, we assessed their performance segmenting the HC and surrounding temporal horns of the lateral ventricles. The measures of performance were: 1) volumetric overlap between segmentations from the automatic methods and the manual segmentation calculated using Dice’s Kappa metric, and 2) Spearman’s rank correlation coefficients between computed volumes from the automatic methods and volumes from the manual segmentations. Note Spearman’s rank correlation was used because the volume data was not Gaussian distributed, evidenced by a significant Shapiro-Wilk test.

We compared the difference in overlap with manual labels for each side of all segmentation methods using Kruskal-Wallis one-way analysis of variance by ranks. We then did post-hoc analyses on the significant results using the Dunn’s test for multiple comparisons with Bonferroni correction.

#### 2.5.2. Application on clinical data

We evaluated the different automatic segmentation techniques by applying them to the baseline data from ADNI. The first comparison was the percentage of failure for the different methods. An expert visually inspected the output of the trained automatic methods and discarded any segmentation that did not pass quality control. For the HC volumes from FreeSurfer versions 4.3 & 5.1, we assumed failure for any missing value on the UCSF volumes files reported by ADNI.

We then compared the different methods by evaluating for both HCvol and HVR: 1) the similarity across segmentations, and 2) the sensitivity of the different methods in detecting between-group differences. To measure the similarity between methods we used the Spearman’s rank correlation coefficient between the computed hippocampal measures—stratified by clinical label. For the sensitivity comparison, we computed the standardized effect size estimates and their robust confidence intervals (CIs) of the difference between the hippocampal measures between the patients with dementia and the cognitively normal subjects. We used Glass’ 𝛥, where the comparison of the means is standardized only by the control’s group standard deviation:

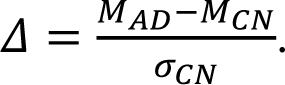

The CIs were calculated using the bias-corrected-and-accelerated bootstrap method (Efron 1987) with the default setting of 2,000 resamples.

### 2.6. Associations of hippocampal volumetrics with age, memory, and global cognition

We also measured the associations of the HCvol and HVR obtained from the automatic segmentation methods with age, memory and global cognition using Spearman’s rank correlation coefficients, given the non-Gaussian distribution of the data. To test the hypothesis that HVR would better associate with these covariates than HCvol, we used single-tailed permutation tests on the difference between the correlation coefficients of these two hippocampal measures. The permutation tests were run with 10,000 repetitions each. The p-values for the correlations and the permutation tests were adjusted using the Bonferroni correction for multiple comparisons.

### 2.7. Data, materials, and software availability

MRI, neuropsychological, and demographic data are available from the ADNI database (https://adni.loni.usc.edu). All of the statistical analyses were done using R version 4.2.2. We used the BootES package (Kirby and Gerlanc 2013) to calculate the effect size estimates and their CIs. All code required to reproduce the analyses is available on GitHub: https://github.com/soffiafdz/hvr_validation.

## 3. Results

Our sample consisted of 1,641 participants in ADNI-1, ADNI-Go and ADNI 2: 501 cognitively healthy participants (CH), 819 individuals with MCI and 321 patients with clinically probable AD. All demographic and clinical measures were significantly different among the three groups (Table 1). The Chi-squared test showed significant differences of sex for the CH (p = 0.002) and MCI (p = 0.004) groups, but not for AD (p = 1). The post-hoc analysis using Dunn’s test showed that participants in the MCI group were significantly younger than those of the CH group (Z = 2.6, p = 0.014) and the AD group (Z = 4.03, p < 0.001). However, the age difference between the CH and AD groups was non-significant (Z = 1.65, p = 0.15). The CH group also appeared to be on average slightly more educated compared to both the MCI group (Z = 2.64, p = 0.012), and the AD group (Z = -5.59, p < 0.001). The difference between the MCI and AD groups was also significant (Z = -3.8, p < 0.001).

**Table 1:** Demographic data

| Variable | N | Clinical Label |  |  | p-value<sup>2</sup> |
| --- | --- | --- | --- | --- | --- |
|  |  | CH, N = 501<sup>1</sup> | MCI, N = 819<sup>1</sup> | AD, N = 321<sup>1</sup> |  |
| Sex | 1,641 |  |  |  | <0.001 |
| Female |  | 260 (52%) | 336 (41%) | 146 (45%) |  |
| Male |  | 241 (48%) | 483 (59%) | 175 (55%) |  |
| Age (years) | 1,641 | 74 (6) | 73 (8) | 75 (8) | <0.001 |
| Education (years) | 1,641 | 16 (3) | 16 (3) | 15 (3) | <0.001 |
| ADAS13 | 1,619 | 9 (4) | 17 (7) | 30 (8) | <0.001 |
| Missing |  | 2 | 9 | 11 |  |
| RAVLT (learning) | 1,629 | 5.85 (2.30) | 4.07 (2.59) | 1.81 (1.79) | <0.001 |
| Missing |  | 4 | 4 | 4 |  |

<sup>1</sup>n (%); Mean (SD)
<sup>2</sup>Pearson's Chi-squared test; Kruskal-Wallis rank sum test

Table 1. Sex, age, education, and the scores for ADAS13 and the learning subtest of the RAVLT for the CH, MCI and AD groups of the baseline data obtained from the ADNI dataset.

### 3.1. Training of the automatic segmentation methods

We trained three automatic segmentation methods—MALF, NLPB, and CNN—using a set of manually segmented labels of HC and CSF. Figure 1 shows the results of this training in the form of violin plots of the volumetric overlap with the manual labels and one example subject’s manual and computed labels from all three methods.

**Figure 1.**
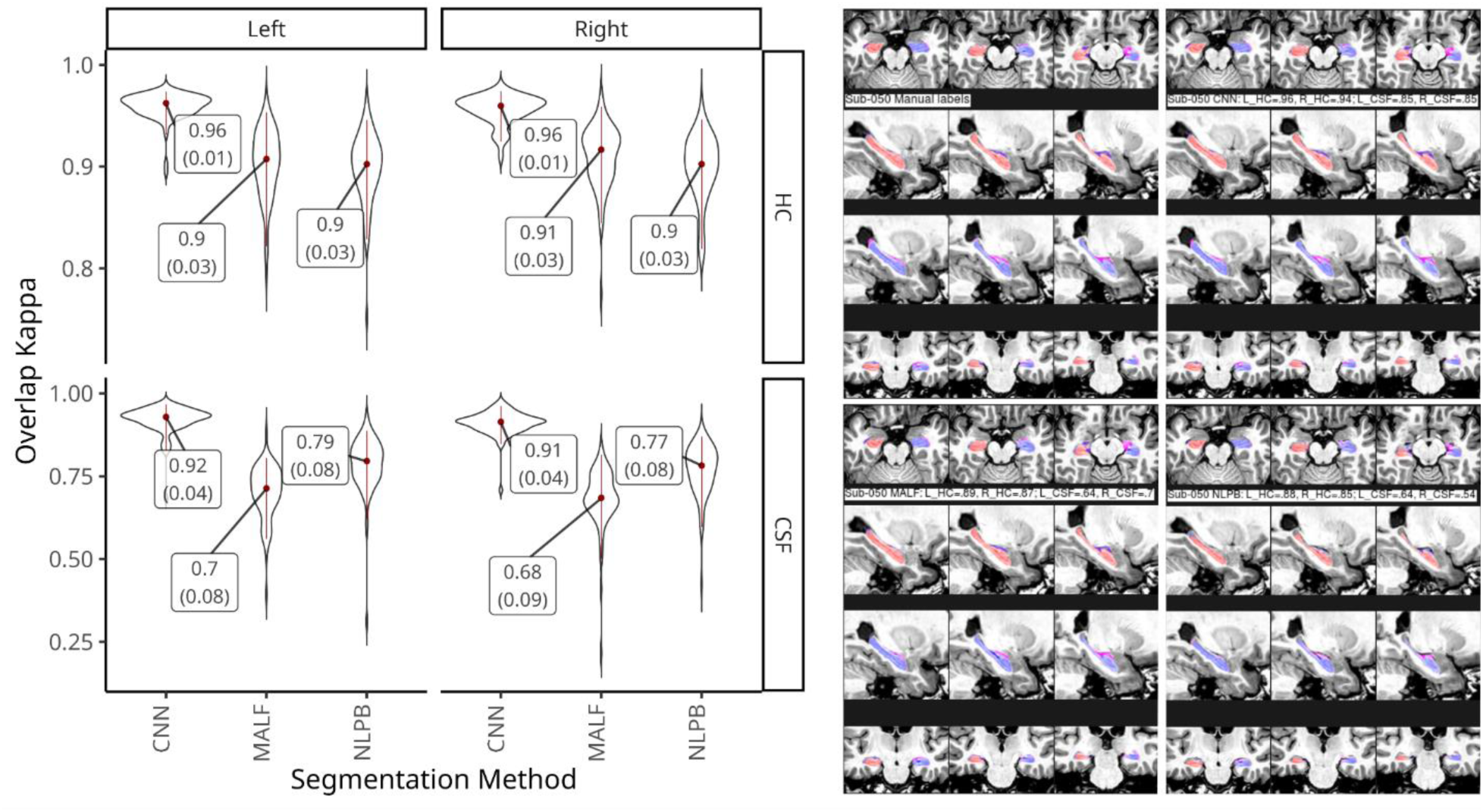
Automatic segmentation performance. Left: violin plots of the Kappa’s distributions of the overlap between manual and computed segmentations of the HC and CSF from the left and right hemispheres using CNN, MALF and NLPB. The median and standard deviation are represented in red inside the violin plots and marked as labels. Right: Mosaic images of the manual and computed segmentations of the HC and CSF for an example subject.

The CNN-based method showed the best performance in both HC and CSF segmentations with a median Kappa value K = 0.96 for both HC sides, with K = 0.92 and K = 0.91 for left and right CSF in the temporal horns of the lateral ventricles, respectively. The Kruskal-Wallis test of Kappa values showed significant differences between the different segmentation methods for the HC (left: X2 = 141.32, p < 0.001; right: X2 = 132.06, p < 0.001) and CSF regions (left: X2 = 167.4, p < 0.001; right: X2 = 173.04, p < 0.001). Post-hoc analyses showed that, for both sides, CNN was better than NLPB and MALF in segmenting HC (p < 0.001), while the performance of NLPB and MALF were comparable (left: p = 0.892; right: p = 0.14). When segmenting CSF, all comparisons between the three methods were statistically significant (p < 0.001).

Figure 2 shows correlations between volumes obtained from the original manual labels and those from computed segmentations. All associations were significant (p < 0.001). The Spearman’s rho values for the CNN method were almost 1 for both HCvol and CSFvol on both left and right sides. While the correlations were lower for NLPB, they still showed strong associations with the manual labels (HCvol, rho = 0.93; CSFvol, rho = 0.85). MALF showed the lowest correlations among the three methods (HCvol, rho = 0.87; CSFvol, rho = 0.48).

**Figure 2.**
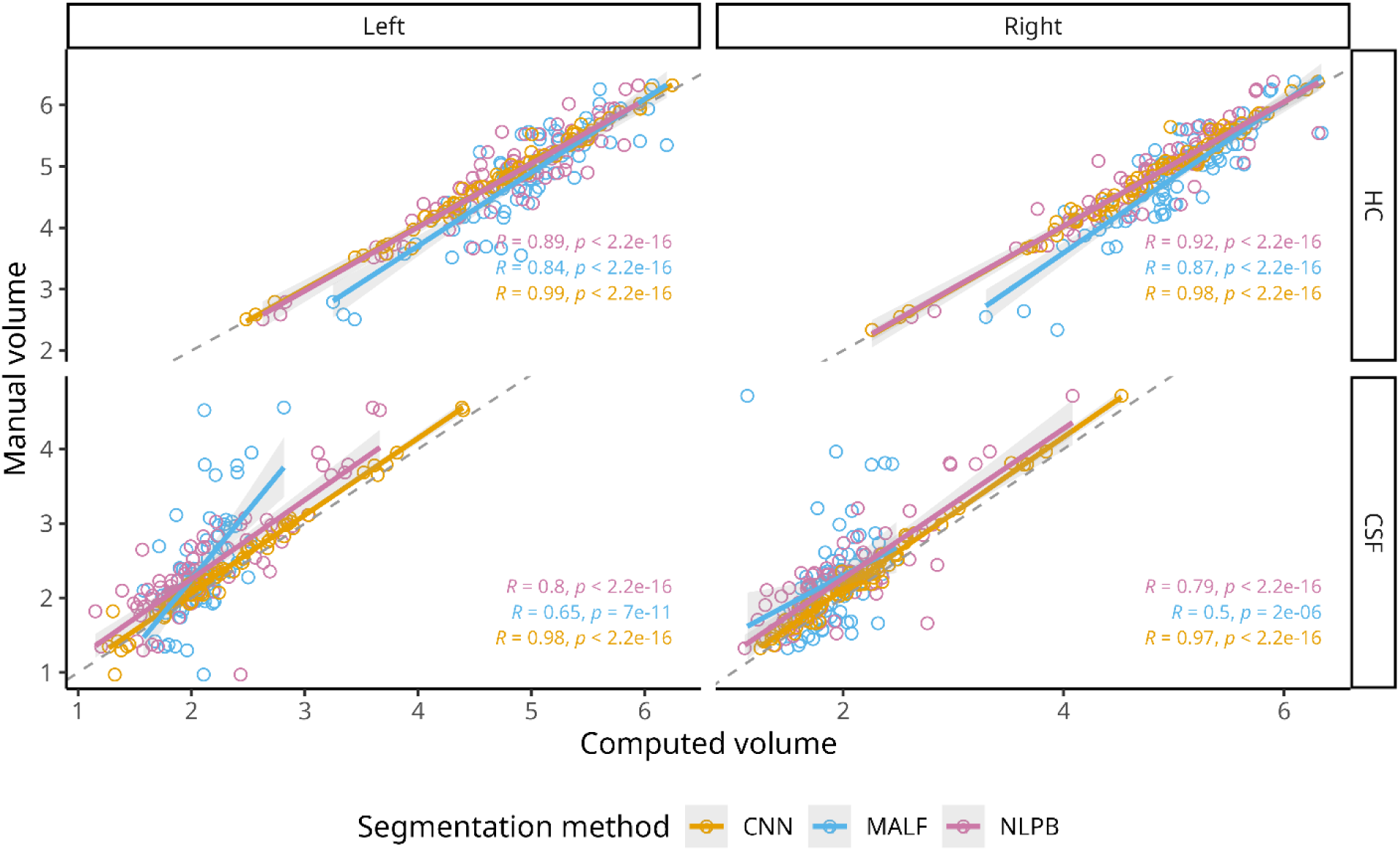
Relationship between manual and computed values of HCvol and CSFvol. Scatter plots of the volumes (measured in cc) of the left and right HC and CSF from the manual segmentations against those obtained from CNN (orange), MALF (blue) and NLPB methods (green). The Spearman’s rho coefficients and their significance are added for each automatic method.

The Bland-Altman plots in Fig. 3 show that in addition to having larger error, MALF had a systematic bias of over-segmenting smaller HC and under-segmenting the larger HC. While NLPB had a moderate variance in error, we did not find a significant pattern of bias in its segmentations. All three methods tended to underestimate CSFvol on average, with MALF also showing a bias of under-segmenting larger ventricles compared to the smaller ones.

**Figure 3.**
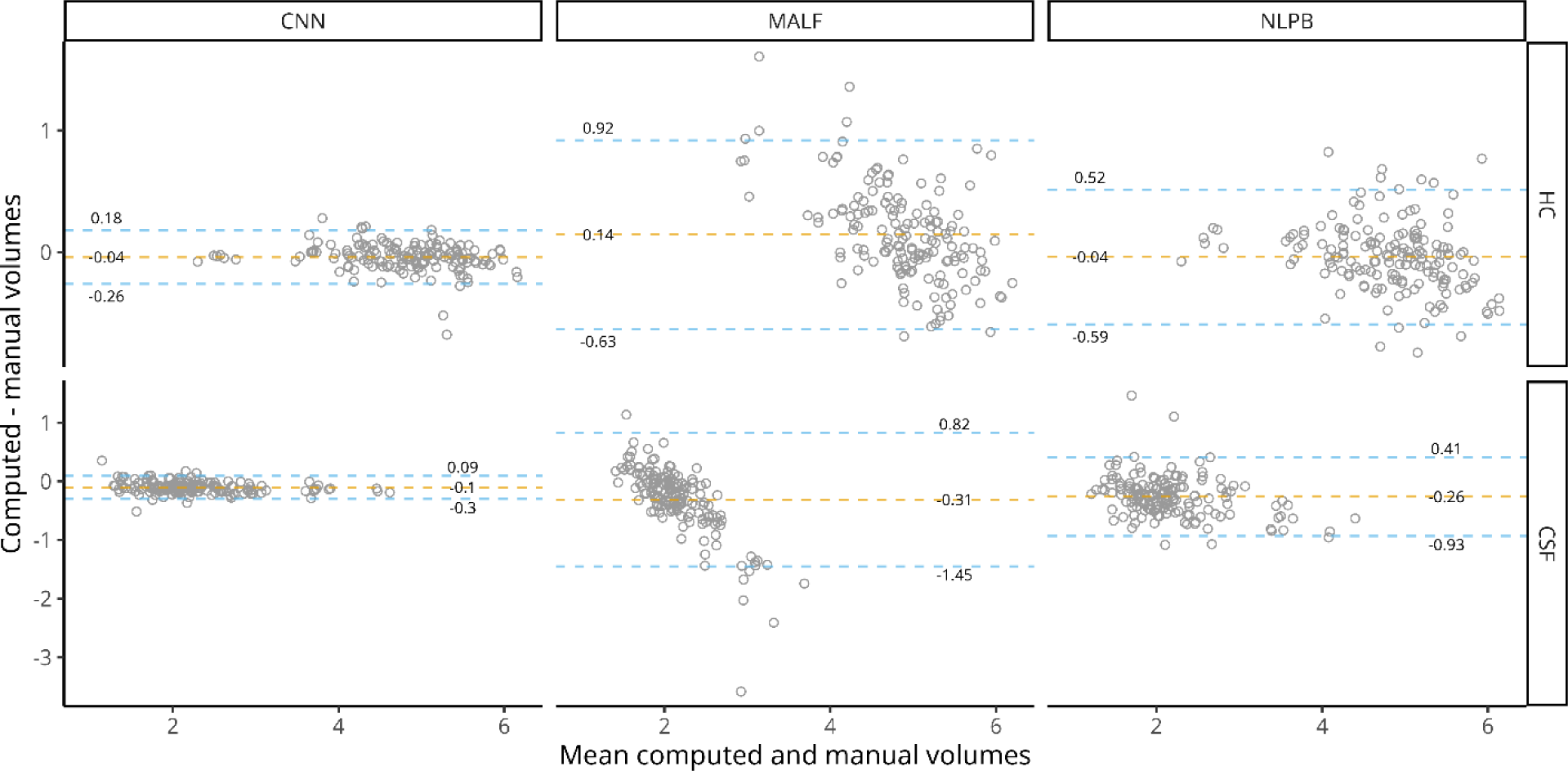
Bland-Altman plots. Bland-Altman plots for the volumetric comparison between mean HCvol and CSFvol (in cc) resulting from the automatic segmentation methods and manual labels. The dashed lines represent the mean difference (orange) and the upper and lower limits of agreement (blue). All volumes are reported in cubic centimeters.

### 3.2. Application on clinical data

We applied the three trained automatic segmentation methods and compared them to the values obtained using the widely used FreeSurfer labels. On Table 2 we report the number of successful HC segmentations for the different methods and the mean HCvol for each of the clinical groups.

**Table 2:** HCvols and segmentation failures

| Automatic Method | N | Clinical Label |  |  |
| --- | --- | --- | --- | --- |
|  |  | CH, N = 501<sup>1</sup> | MCI, N = 819<sup>1</sup> | AD, N = 321<sup>1</sup> |
| CNN | 1,639 | 4.18 (0.55) | 3.73 (0.71) | 3.16 (0.65) |
| Failures |  | 0 | 1 | 1 |
| NLPB | 1,627 | 4.14 (0.49) | 3.79 (0.61) | 3.38 (0.53) |
| Failures |  | 2 | 9 | 3 |
| MALF | 1,617 | 4.37 (0.39) | 4.19 (0.42) | 4.06 (0.41) |
| Failures |  | 3 | 8 | 13 |

Table 2. Summary of HCvols and segmentation failures.
| **Automatic Method** | **N** | **Clinical Label** |  |  |
| --- | --- | --- | --- | --- |
|  |  | **CH, N = 501<sup>1</sup>** | **MCI, N = 819<sup>1</sup>** | **AD, N = 321<sup>1</sup>** |
| FreeSurfer (v4.3 & v5.1) | 1,414 | 5.16 (0.71) | 4.63 (0.92) | 3.90 (0.72) |
| Failures |  | 47 | 122 | 58 |
| FreeSurfer (v6.0) | 1,616 | 5.04 (0.68) | 4.50 (0.90) | 3.85 (0.71) |
| Failures |  | 8 | 9 | 8 |

<sup>1</sup>Mean (SD)

Table 2. Summary of HCvols and segmentation failures. Mean and standard deviation of the HCvol obtained and the number of failed cases by the different automatic segmentation methods. Note that for v4.3 & v5.1 of FreeSurfer we report the missing data from ADNI rather than explicit failures for the algorithm.

The method with the least number of successful segmentations was FreeSurfer v4.3/v5.1, with ADNI reporting the HC values for only 86% of the data. FreeSurfer v6.0 and MALF had a similar number of successful segmentations (98.5%), nonetheless MALF tended to fail more on AD and MCI participants while FreeSurfer’s errors were equally distributed between all groups. From all methods, CNN was the most reliable, failing on only two cases (one MCI and one AD).

The comparisons between segmentation methods in terms of volumetric correlation and effect size between CH individuals and patients with AD are presented in Fig. 4. HC volumes obtained from the FreeSurfer, CNN and NLPB segmentations were all highly correlated with each other (rho > 0.9, p < 0.001), with the FreeSurfer versions being the most similar between each other (rho = 0.99). The correlations coefficients for MALF, the least similar method, ranged between 0.67-0.78. Even after stratifying the comparisons by clinical groups, we still found high similarity between FreeSurfer versions and CNN (rho > 0.97, between FreeSurfer versions; rho > 0.91, CNN & FreeSurfer). Across all methods, the lowest correlation coefficients were found on the AD group (rho = 0.4 – 0.98).

**Figure 4.**
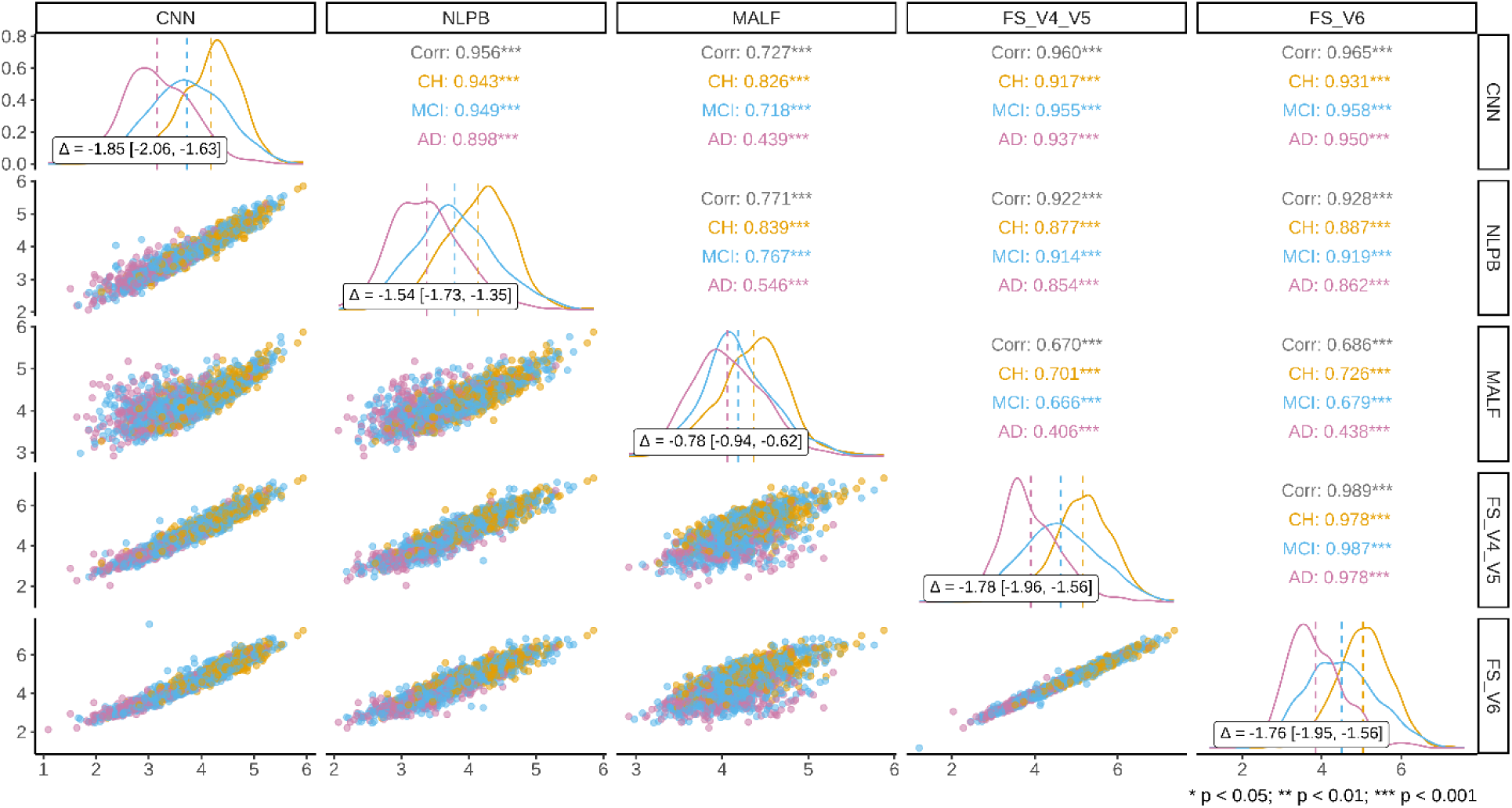
Similarity of segmentation and clinical effect sizes of HCvol. The lower triangle of the matrix plot shows scatter plots of the HCvol values across the different automatic segmentation methods stratified by the clinical group (CH: orange, MCI: blue, and AD: green). The upper triangle of the matrix plots lists the correlation coefficients between the automatic segmentation methods using Spearman’s rho. The correlation coefficients include the values for the whole sample in gray and by clinical group. The diagonal of the matrix plot shows the density plots of the HCvol by clinical group. Dashed lines represent the mean of each cohort. Inside each density plot, the inlaid box shows calculated effect sizes between the CH and AD groups using Glass’ Delta and their robust 95% confidence intervals.

Effect sizes calculated using Glass’ delta follow the conventional interpretation of Cohen’s d small, medium, and large effect sizes defined as d < 0.5, 0.5 < d < 0.8, and d > 0.8, respectively. We found large effect sizes on the HCvol difference between CH and AD groups for all segmentation methods except MALF. The methods, from worst to best, were MALF (Δ = -0.78), NLPB (Δ = -1.54), FreeSurfer (Δ = -1.76 – -1.78), and CNN (Δ = -1.85). Comparing the CH:AD effect sizes between the segmentation methods, the only significant differences (based on the calculated confidence intervals) were observed on the ones from MALF which were lower compared to the other three automatic methods.

Using the values for HCvol and CSFvol, we calculated the HVR for all segmentation methods; the mean and standard deviation for the HVR values by clinical group are given in Table 3. HVR values were larger for the FreeSurfer-based estimates. The correlations between methods and the comparison of effect sizes between CH and AD are presented in Fig. 5. The similarity across methods was higher for HVR (compared to HCvol) with the correlation coefficients ranging from 0.89 (MALF & FreeSurfer) to 0.97 (CNN & FreeSurfer). As with HCvol, the lower correlations were found on the AD group (rho = 0.77 – 0.97). All four segmentation methods had large effect sizes between CH and AD when using HVR. The magnitudes of the effect size from HVR were larger than those calculated with HCvol, nonetheless, the HVR vs HCvol difference in effect size only reached significance for MALF. Ordered from lowest to highest value, the methods were MALF (Δ = -1.24), NLPB (Δ =-1.73), CNN (Δ = -1.86), and FreeSurfer (Δ = - 2.02). The effect size obtained with MALF was significantly lower than the other methods. None of the other differences across methods were significant.

**Table 3:** HVR values

| Automatic Method | N | Clinical Label |  |  |
| --- | --- | --- | --- | --- |
|  |  | CH, N = 501<sup>1</sup> | MCI, N = 819<sup>1</sup> | AD, N = 321<sup>1</sup> |
| CNN | 1,639 | 0.64 (0.08) | 0.58 (0.11) | 0.49 (0.11) |
| NLPB | 1,627 | 0.67 (0.08) | 0.62 (0.10) | 0.54 (0.11) |
| MALF | 1,617 | 0.69 (0.03) | 0.67 (0.04) | 0.65 (0.03) |
| FreeSurfer (v6.0) | 1,616 | 0.84 (0.08) | 0.77 (0.12) | 0.67 (0.13) |

<sup>1</sup>Mean (SD)

**Figure 5.**
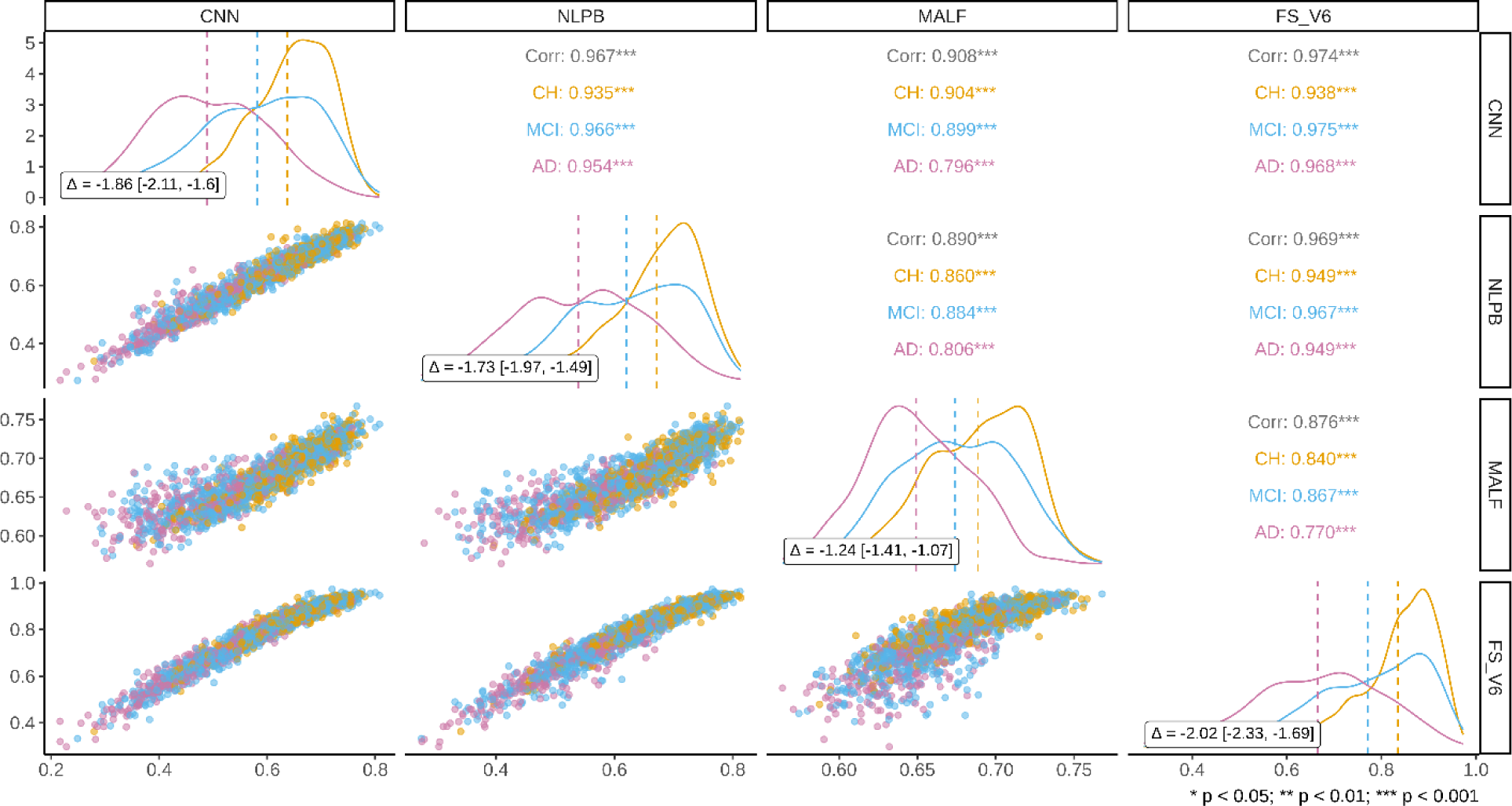
Similarity of values of HVR and their derived clinical effect sizes. The lower triangle of the matrix plot shows scatter plots of the HVR values across the different automatic segmentation methods stratified by the clinical group (CH: orange, MCI: blue, and AD: green). The upper triangle of the matrix plots lists the correlation coefficients between the automatic segmentation methods using Spearman’s rho. The correlation coefficients include the values for the whole sample in gray and by clinical group. The diagonal of the matrix plot shows the density plots of the HVR by clinical group. Dashed lines represent the mean of each cohort.

Table 3. Summary of HVR values. Mean and standard deviation of the HVR values obtained from the different automatic segmentation methods.

Inside each density plot, the inlaid box shows calculated effect sizes between the CH and AD groups using Glass’ Delta and their robust 95% confidence intervals.

### 3.3 Associations of hippocampal volumetrics with age, memory, and global cognition

The correlation coefficients between HCvol with age, the subscores of learning from RAVLT and the scores of ADAS13 for all four segmentation methods are presented in Figure 6. The correlation coefficients for HVR are presented as well. All correlations and comparisons were corrected for multiple comparisons. We found a significant positive association (p < 0.001) between HCvol and Age across all clinical groups for FreeSurfer (N = 1,589), CNN (1,612), and NLPB (N = 1,600); for MALF (1,590), the correlation was significant only on the CH (p = 0.002) and MCI groups (p < 0.001). The association between HVR and age was significant for all groups and methods (p < 0.001). The correlations of age with HVR were higher compared to correlations with HCvol. This difference was significant on the CH and MCI cohorts for the CNN (CH, rho: -0.4 vs -0.53, p = 0.012; MCI, rho: -0.46 vs -0.57, p = 0.01) and NLPB (CH, rho: -0.34 vs -0.49, p = 0.006; MCI, rho: -0.39 vs -0.54, p = 0.001), and all clinical groups of MALF (CH, rho: -0.19 vs -0.5; MCI, rho: -0.2 vs -0.52; AD, rho: -0.03 vs -0.38; all p < 0.001).

**Figure 6.**
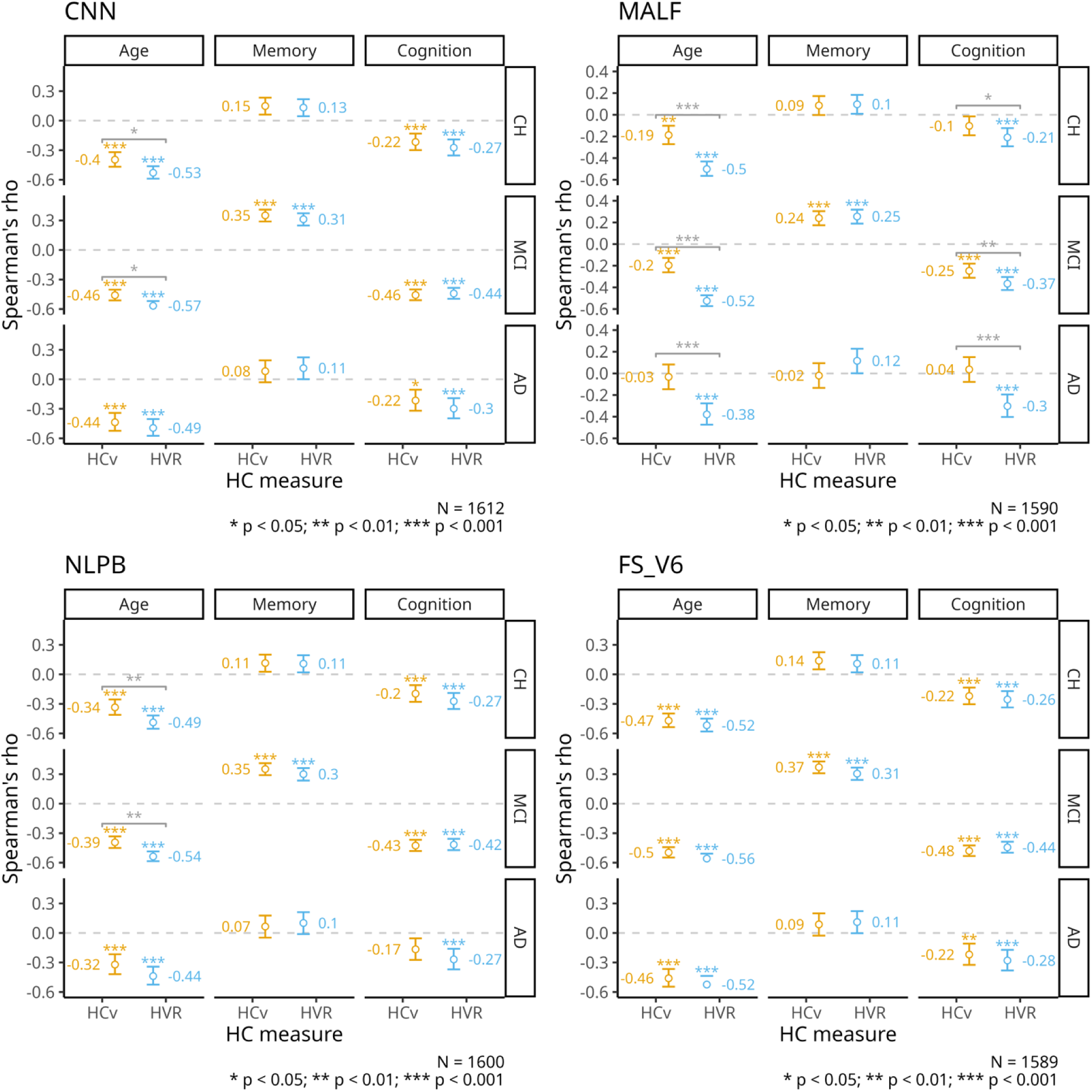
Associations between hippocampal measures and age, memory, and global cognition. Summary of the correlation coefficients measured by Spearman’s rho, between HCvol (HCv in the legend), HVR, and age, memory (measured by RAVLT learning subscore), and global cognition (measured by ADAS13). Colored asterisks mark the significance of the correlation (*= p < 0.05; ** = p <0.01, *** = p < 0.001). Gray asterisks mark the significance comparing HCvol and HVR. All p-values are corrected for multiple comparisons.

There was a positive association with the RAVLT learning subscore on both HCvol and HVR with all four methods, but this association was significant only for the MCI cohort (HCvol: 0.24 – 0.37; HVR: 0.25 – 0.31; p < 0.001). None of the differences in correlations of HCvol and HVR with memory were statistically significant.

Lastly, we noted a negative association between both HCvol and HVR with the ADAS13 scores across groups. The correlations of HCvol with cognitive decline were significant in the CH group segmented with FreeSurfer (rho = -0.22, p < 0.001), CNN (rho = -0.22, p < 0.001) and NLPB (rho = -0.2, p <0.001), but not MALF (rho = -0.1, p = 1); and the MCI group on all four methods (FreeSurfer: rho = -0.48; CNN: rho = -0.46; NLPB: rho = -0.43; and MALF: rho = - 0.25; p < 0.001). For the AD group, the only significant correlations between HCvol and ADAS13 were found on the FreeSurfer (rho = -0.22, p = 0.01) and CNN methods (−0.22, p = 0.01).. The association between HVR and ADAS13 was significant in all three groups of the four segmentation methods (rho = -0.21 – -0.44, p < 0.001). The magnitude of these associations was comparable to those of HCvol on all segmentation methods except for MALF. The correlation between ADAS13 and the HVR obtained from the MALF was stronger on all groups (CH, rho: - 0.1 vs -0.21, p = 0.048; MCI, rho: -0.25 vs -0.37, p = 0.009; AD, rho: 0.04 vs -0.3, p < 0.001).

## 4. Discussion

In this article, our goal was to automate the estimation of the HVR, a robust measure of hippocampal integrity (Schoemaker et al. 2019), to extend the preliminary evidence of its validity as a measure of hippocampal atrophy in a much larger clinical sample.

To fulfill our goal, we evaluated the viability of various automatic segmentations to calculate the HVR. Ensuring a fair assessment of performance when comparing different automatic segmentation methods is not a trivial task. Segmentation accuracy is affected by several factors, including the quality of the MRI data, the segmentation protocol used for the ground-truth, the particularities of the sample tested, and the metric used to evaluate the performance (Collins and Pruessner 2010). A practical example of this issue is the use of Dice Kappa as a measurement for segmentation accuracy. Given the metrics dependance on the surface to volume ratio, Dice Kappa measurements are stricter on smaller regions. This can be significant when evaluating the automatic segmentations that comprise the HVR: the hippocampus, a small and highly variable structure (Lupien et al. 2007; Nobis et al. 2019), and the even smaller surrounding ventricular space.

Despite this inherent drawback, all three of the evaluated locally developed methods could successfully segment the hippocampus with high accuracy, with the CNN (U-Net based) method showing a clear advantage over the other two. Still, the MALF and NLPB segmentations results were comparable to the accuracy reported in their original papers (Collins and Pruessner 2010; Coupé et al. 2011; Fonov et al. 2012), and even similar to some U-net based methods (Brusini et al. 2020; Cao et al. 2018; Goubran et al. 2020; Liu et al. 2020; Yao, Wang, and Fu 2019). We did not estimate Dice Kappa with Freesurfer as it uses a different anatomical definition for the HC and CSF spaces.

The segmentation of the temporal horn of the lateral ventricle, being a much smaller region with somewhat arbitrary limits in its anterior and posterior borders, was a significantly harder task compared to the extraction of the hippocampus. This increased difficulty was notable in the lower Dice Kappa values compared to the hippocampal segmentation across all methods. However, when measuring the segmentation quality by the volumetric correlation, the results for both CSF and HC segmentation tasks were comparable to the CNN method.

The Bland-Altman plots in Figure 3 confirmed the superior performance of the CNN technique, highlighting minimal errors and a lack of bias. This stands in contrast to the wider limits of error observed in NLPB and the bias exhibited by MALF in the form of a regression to the mean.

When applied to the baseline data of ADNI, the CNN method demonstrated superior reliability amongst all segmentation methods, with less than 0.2% failed segmentations. We found the earliest versions of FreeSurfer to be the least dependable method of our comparisons, given the absence of ∼14% of the values for hippocampal volume reported by ADNI. It is important to note that the assumption that all missing values are due to segmentation failures may be unfair. In addition, the FreeSurfer team likely uses QC criteria different than ours. Nonetheless, this assumed failure rate would align with a recent QC evaluation of the output of a more recent version of FreeSurfer (v6.0), on which 5% of the segmentations were manually marked as failures and 20% were marked as having doubtful quality (Klapwijk et al. 2019). Notably, in our own use, FreeSurfer v6.0 had a comparable failure rate to the other label-fusion based methods.

Despite expected improved results from the updated release of FreeSurfer, the hippocampal segmentations were not drastically different as our implementation of v6.0 yielded segmentations that were highly correlated to the UCSF volumes reported by ADNI. For both versions, FreeSurfer tended to yield higher HCvol values compared to the three other methods evaluated. This difference is not surprising. First, the training library used for FreeSurfer was not the same as that of the template library of Schoemaker (2019) used in our training. Second, the underlying anatomical definition of the hippocampus is different between FreeSurfer and the labels of Schoemaker. Additionally, the tendency of FreeSurfer to over segment larger hippocampus has already been noted in the literature (Pipitone et al. 2014; Zandifar et al. 2017).

Regardless of this over segmentation of HC from FreeSurfer, its volumetric measurements were highly correlated to those obtained from the CNN method and, to a slightly lesser extent, from the NLPB method. These three methods also had larger effect sizes between healthy and AD cohorts that were comparable to each other and significantly different to the medium effect size obtained from the underperforming MALF.

HVR was proposed as a more robust measure of hippocampal atrophy compared to HCvol. Our experiments showed increased similarity of the HVR calculation compared to HCvol among the different segmentation techniques. Furthermore, the CH:AD HVR effect sizes were larger than CH:AD HCvol effect sizes for all methods tested (Figs 4 & 5). This improvement was particularly more evident on the previously underperforming MALF method, for which the similarity to the other algorithms increased by a magnitude of ∼ 0.2 in rho and the CH:AD effect size improved from 0.78 to 1.24 (a 57% improvement) in the Glass’ delta. It would seem that HVR is more useful for poorer quality segmentations.

FreeSurfer’s combination of some over segmentation of larger hippocampi with the use of only the inferior portion of the temporal horn of the lateral ventricle resulted in much higher HVR values compared to the three other methods. These greater values led to a more pronounced separation of healthy and atrophied HC, which resulted in the largest effect size of 2.02 between CH and AD across all methods.

There is an expected hippocampal atrophy due to aging even in healthy individuals (Bettio, Rajendran, and Gil-Mohapel 2017). The age-related declines in hippocampal volume found in our healthy cohort (Fig. 6) were in the upper ranges of what has been found in previous reports (for reviews see: (Raz 2000; Van Petten 2004). The rho values for HVR tended to be stronger compared to HCvol across all cohorts and all methods, except for FreeSurfer, as FreeSurfer displayed strong correlations for HCvol with age already. Incidentally, for HCvol, FreeSurfer’s tendency to over-segment larger hippocampi and under-segment smaller ones might have played a role in the higher sensitivity of this method in capturing the effect of aging, as the magnitude of this relationship even surpassed HVR’s association with age reported in the original Schoemaker paper (2019).

While this is the first time that HVR is applied to a clinical sample, the study of hippocampal atrophy in MCI and AD compared to normal aging has been heavily studied over the years (den Heijer et al. 2010; Ikram et al. 2010; Jack et al. 1997; Pol et al. 2006), noting smaller hippocampal volumes compared to controls (Frankó, Joly, and Alzheimer’s Disease Neuroimaging Initiative 2013; Jack et al. 1997; 2000; Morra et al. 2009), and even the potential of using hippocampal rates of decline to differentiate stable and progressive MCI (Ikram et al. 2010; P.-N. Wang et al. 2009). These results coincide with our slightly stronger associations with age in the MCI cohort compared to the healthy subjects for both hippocampal measures, again with a significant advantage of HVR for all segmentations methods except FreeSurfer. Meanwhile, for the AD patients, HVR showed a statistical advantage only for MALF which, in addition to having the lowest rho values overall, was the only method not sensitive enough to find a significant association between HCvol and age in this cohort, likely because the MALF estimated volumes regress to the mean.

An assumed relationship between hippocampal volume and memory exists, with larger hippocampi associated with better scores on neuropsychological testing. We found such a relation in our MCI subgroup for all segmentation methods, but not in the cognitively normal or AD groups. These results contrasted with previous reported relationships in AD between the delayed subtest of RAVLT with HOC in a Brazilian sample (Sudo et al. 2019) and with HCvol within ADNI (X. Wang et al. 2019).

Further, HVR associations with memory were comparable to those of HCvol among all segmentations and subgroups. While these results contrast with the preliminary report of HVR surpassing an already high association between HCvol and memory measured by the same test, they fit with the heterogeneous and highly variable existing evidence for the relationship between HCvol and memory performance (Van Petten 2004). Another potential source of difference between our results and those reported by Schoemaker (2019) is that our groups were on average a decade older, and greater individual variability in episodic memory has been found for older adults (Morse 1993; Nelson and Dannefer 1992; Verhaeghen and Marcoen 1990; Wilson et al. 2002). Additionally, given the clinical nature of our sample, we also explored the association between our hippocampal measures and a more global evaluation of cognition in the form of the ADAS13, which, in addition to memory, assesses other domains like language and praxis (Kueper, Speechley, and Montero-Odasso 2018). Previous studies have found a relationship between the hippocampus and global cognitive functioning measured with the mini-mental status examination (MMSE) in only AD (Vipin et al. 2018), or AD and MCI but not normal controls (Peng et al. 2015). Despite these results and the fact that ADAS13 being designed to evaluate cognitive dysfunction in early stages of AD, we did find a significant association with HCvol in our healthy cohorts. For cognitively normal and MCI subjects, the associations with global cognition of both HVR and HCvol were comparable. Contrary to the abundant evidence of the relationship between hippocampus and general cognition in AD (Morrison, Dadar, Shafiee, and Collins 2023; Morrison, Dadar, Shafiee, Collins, et al. 2023; Peng et al. 2015; Vipin et al. 2018), the correlations in our AD cohort were lower compared to the other subgroups, with only HVR’s reaching statistical significance across all segmentation methods. This advantage of HVR over HCvol was statistically significant for the MALF segmentations of the MCI and AD cohorts, further suggesting that HVR might be particularly superior when only lower quality segmentations are available.

Our study is not without limitations. First, although this is the first time that HVR has been applied to a sample of this magnitude, by using the ADNI data which consists mostly of highly educated white people, our results might not generalize to other more diverse populations. We used only the baseline data available, and future work will explore longitudinal changes in hippocampal degradation. In addition, we did not look into the effects of laterality or sex, even though there were substantial sex differences reported in the original paper on HVR (Schoemaker et al., 2019). This omission was deliberate to avoid increasing the already large number of statistical comparisons. Future research could further validate the HVR measure by evaluating these potential sources of variability. Another limitation is related to the choice of segmentation methods we evaluated. Given the wide use of the FreeSurfer pipeline, we decided to include it in our selection of evaluated methods even though it was impossible to evaluate its accuracy without its gold standards, and unfair to compare it to our gold standards. Relatedly, its comparison of calculated HVR values to our trained methods was tainted by different anatomical definitions of the hippocampus and the surrounding ventricles compared to the other methods. Despite these differences, the high correlation to our proposed CNN method together with the comparable CH:AD effect size and associations with age and global cognition support its use as a viable alternative to our proposed CNN method.

### 4.1 Conclusions

In conclusion, after evaluating various automatic segmentation methods, with Dice Kappas of 0.96 and rho values of 0.99 against high quality manually segmented labels, the CNN method yields the best HC segmentations among the three methods, with results that are comparable to the best in the literature. The high HCvol correlations between FreeSurfer, CNN and NLPB methods show that these methods agree fairly well for the segmentation of the HC, and thus all are viable choices to compute the HVR in large samples. HVR, when compared against HCvol, showed greater consistency across segmentation methods, a larger effect size when distinguishing between individuals with AD and healthy controls, and stronger associations with age and global cognition, particularly improving the performance of the MALF method. These findings support the promise of HVR as a measure of hippocampal integrity, specifically for neurodegenerative disease research and the study of age-related cognitive decline.

## Acknowledgements

Data collection and sharing for this project was funded by the Alzheimer’s Disease Neuroimaging Initiative (ADNI) (National Institutes of Health Grant U01 AG024904) and DOD ADNI (Department of Defense award number W81XWH-12-2-0012). ADNI is funded by the National Institute on Aging, the National Institute of Biomedical Imaging and Bioengineering, and through generous contributions from the following: AbbVie, Alzheimer’s Association; Alzheimer’s Drug Discovery Foundation; Araclon Biotech; BioClinica, Inc.; Biogen; Bristol-Myers Squibb Company; CereSpir, Inc.; Cogstate; Eisai Inc.; Elan Pharmaceuticals, Inc.; Eli Lilly and Company; EuroImmun; F. Hoffmann-La Roche Ltd and its affiliated company Genentech, Inc.; Fujirebio; GE Healthcare; IXICO Ltd.; Janssen Alzheimer Immunotherapy Research & Development, LLC.; Johnson & Johnson Pharmaceutical Research & Development LLC.; Lumosity; Lundbeck; Merck & Co., Inc.; Meso Scale Diagnostics, LLC.; NeuroRx Research; Neurotrack Technologies; Novartis Pharmaceuticals Corporation; Pfizer Inc.; Piramal Imaging; Servier; Takeda Pharmaceutical Company; and Transition Therapeutics. The Canadian Institutes of Health Research is providing funds to support ADNI clinical sites in Canada. Private sector contributions are facilitated by the Foundation for the National Institutes of Health (www.fnih.org). The grantee organization is the Northern California Institute for Research and Education, and the study is coordinated by the Alzheimer’s Therapeutic Research Institute at the University of Southern California. ADNI data are disseminated by the Laboratory for Neuro Imaging at the University of Southern California.

## Abbreviations

AD: Alzheimer’s disease
ADAS13: Alzheimer’s disease assessment scale 13
ADNI: Alzheimer’s disease neuroimaging initiative
CI: confidence interval
CH: cognitively healthy
CSF: cerebro-spinal fluid
CSFvol: perihippocampal ventricular space
CNN: convolutional neural network
HC: hippocampus
HCvol: hippocampal volume
HVR: hippocampal-to-ventricle ratio
MALF: multi-atlas label fusion
MCI: mild cognitive impairment
MRI: magnetic resonance imaging
NLPB: non-local patch-based
RAVLT: Rey auditory verbal learning test.

## Notes

### Competing Interest Statement

The authors have declared no competing interest.

